# Dorsal Striatum Silencing Attenuates Light Self-administration in Mice and Its Relevance to Digital technology-based Disorders

**DOI:** 10.64898/2026.07.31.742007

**Authors:** Shu K. E. Tam, Benjamin Becker

## Abstract

**Background and aims:** Substance-use addiction models have demonstrated that a progressive transition from reward-guided to habitual and ultimately compulsive behaviour is mediated by a ventral-to-dorsal striatal shift in behavioural control. While symptomatic and neural similarities between substance and digital technology-based disorders have been hypothesised, the causal role of the striatum in the latter remains unknown. We here employ a validated mouse light self-administration paradigm with a loss-of-function approach to determine the role of the dorsal striatum in behavioural persistence towards non-food, non-drug reinforcers.

**Methods:** Mice received a control virus or a virus expressing the inhibitory designer receptor (hM4Di) into the dorsal striatum (caudate–putamen). The chemogenetic actuator clozapine N-oxide (CNO) was injected systemically shortly before selected sessions, silencing striatal neuronal firing *in vivo* in hM4Di-expressing mice. During operant training, lever presses were reinforced by light under fixed-ratio schedules of reinforcement (FR1, FR3, and FR5).

**Results:** On initial training days, CNO reduced light self-administration in striatal hM4Di-expressing mice. CNO had a negligible effect in control mice under FR3 and FR5 but attenuated responding in hM4Di mice, which showed response recovery on days without CNO. Linear mixed-effects models confirmed an improvement in light self-administration across days, with a stronger detrimental effect of CNO in hM4Di mice.

**Discussion:** Dorsal striatum silencing attenuates light self-administration without impairing response acquisition. Thus, like drug self-administration, light self-administration relies partly on dorsal striatal neurons. Our work bridges the gap between animal models and human neuroimaging studies reporting shared brain mechanisms underlying non-drug and drug habits.

## Introduction

Digital technology-based disorders have become a growing concern in an increasingly connected society (Király, Koncz, Griffiths, & Demetrovics, 2023; Montag & Becker, 2023). The dysregulated use of digital services has been increasingly conceptualised as a behavioural addiction, with excessive online gaming being recognised as (Internet) gaming disorder in the *ICD-11* (World Health Organization, 2026) and as a condition warranting further study in the *DSM-5* (American Psychiatric Association, 2022), reflecting a hypothesised overlap of symptomatic and neural features with substance use disorders (Brand et al., 2025) and other forms of compulsion-related disorders (Robbins, Banca, & Belin, 2024; Klugah-Brown et al., 2021). Virtual reward stimuli and processes that may drive initial excessive engagement with digital devices are multifaceted, encompassing multiple sources of content-derived reinforcement; which include in addition to sensory rewarding experience also information seeking, cognitive stimulation, boredom relief, and social validation (da Silva Pinho, Céspedes Izquierdo, Lindström, & van den Bos, 2024; Lindström et al., 2021; Montag, Yang, & Elhai, 2021). Rewarding properties of these reinforcers are influenced by individual differences, cognitive states, as well as linguistic, social, and cultural factors, making it challenging to isolate and study the underlying neurofunctional mechanisms in humans (Tam & Becker, 2026).

Animal models examining escalating substance use as a model of human compulsive drug use indicate that at the neural level the initial rewarding experience and progressively escalating use is mediated via the nucleus accumbens in the ventral striatum (Everitt & Robbins, 2005, 2013), a brain region involved in processing information about different primary reinforcers and conditioned incentives (Daniel & Pollmann, 2014). During the transition towards more excessive and ultimately addictive stages of use patterns, the behaviour becomes less reward driven and less regulated by the ventral striatum but progressively controlled by the by the dorsal striatum, a region supporting habits driven by *stimulus* → *response* associations (Everitt & Robbins, 2005, 2013; Yin & Knowlton, 2006). Several human neuroimaging studies reported such a shift from ventral to dorsal striatal circuits in participants with substance dependence (Vollstädt-Klein et al., 2010; Zhou, F. et al., 2018; Zhou, X. et al., 2019) and (Internet) gaming disorder (Dong et al., 2021). These studies provide support for the theoretical viewpoint emphasising the progression from outcome-sensitive reward-seeking behaviour (which depends on the ventral striatum) to outcome-*in*sensitive habits and compulsion (which depends on the dorsal striatum; Everitt & Robbins, 2005, 2013). While this transdiagnostic neural framework of compulsive behaviour has been established for substance use disorders based on sophisticated animal models (Robbins et al., 2024) and is highly relevant to digital technology-based disorders (Klugah-Brown et al., 2021), the underlying circuits have not been directly tested for behavioural addictions.

Although neuroimaging studies can shed light on the functional and structural neural correlates of (excessive) digital engagement (Meshi, Morawetz, & Heekeren, 2013; Montag & Becker, 2023; Montag et al., 2017), they cannot directly confirm the necessity of the dorsal striatum for these conditions; and combining animal models (Casile et al., 2025, 2026) with a loss-of-function approach (e.g., dorsal striatal dysfunction; Jing et al., 2021; Muskens, Schellekens, de Leeuw, Tendolkar, & Hepark, 2012) is critical for this purpose. But unlike substance use disorders where the addictive substance is the primary reinforcer, developing animal models relevant to digital technology-based disorders has been challenging, e.g., due to the multitude and complexity of reinforcers involved (Tam & Becker, 2026). In this regard, sensory reinforcement by LED light may represent a viable option to establish corresponding animal models: sensory and virtual rewards are primarily delivered via LED screens, and there is emerging evidence that LED light at an optimal intensity is intrinsically rewarding in both humans and rodents (Tam, Stryjska, Gu, & Becker, 2025). An advantage of light reinforcers over other non-food, non-drug reinforcers in mice (e.g., thermal cues, novel objects, running wheels, and social interactions with conspecifics) is that the dose, timing, and schedule of light delivery can be precisely controlled, making it well suited for experimental induction of reward-associated behaviour, including behavioural patterns resembling persistent responding despite reduced positive consequences (see e.g., Tam, Xiao, Cheng, Kwok, & Becker, 2026, Exp.3 reinforcer degradation test). Combined with neuronal silencing approaches (e.g., Roth, 2016), this paradigm allows to examine the necessity of the dorsal striatum for the development of escalating sensory self-administration.

In laboratory mice, light self-administration is similar to drug (and food self-administration) with respect to: (*a*) the gradual time course of both response acquisition and response decrement following reinforcer degradation (i.e., time is needed to strengthen and change a habit; Tam et al., 2026, Exp.1 and Exp.3); and (*b*) the necessity of *response* → *reinforcer* contingency, without which light self-administration cannot be established (Tam et al., 2026, Exp.2). Given the involvement of dorsal striatal circuits in (compulsive) drug self-administration in animals (Everitt & Robbins, 2005, 2013; Yin & Knowlton, 2006) and potentially also in humans with substance dependence (Vollstädt-Klein et al., 2010; Zhou, F. et al., 2018; Zhou, X. et al., 2019), we report here an initial investigation on the contribution of the dorsal striatum to light self-administration in laboratory mice by using chemogenetics to temporarily silence dorsal striatal neurons expressing the modified M4 muscarinic designer receptor (hM4Di; Roth, 2016). This allows us to understand overlapping brain regions involved in light and drug self-administration in rodents, revealing shared neural mechanisms underlying non-drug and drug habits in humans.

## Methods

### Animals, housing condition, and ethics

Sixteen naïve 10-week-old male C57BL/6J mice (Model Organisms, Shanghai, China) without any prior operant training experience were used in this study. Each mouse was kept individually with *ad libitum* access to food and water. Animals were housed under a 12-h light:12-h dark cycle, with Zeitgeber time 0 (ZT00) set to 06:00 and ZT12 set to 18:00. Each cage contained a layer of corn cob bedding along with cotton balls for nesting; bedding was replaced every three weeks, and water bottles were changed every ten days. Cages were housed in enclosed wooden chambers with passive-infrared (PIR) sensors (Brown, Hasan, Foster, & Peirson, 2016) mounted 35 cm overhead to monitor daily locomotor activity. During the light phase, mice were exposed to 100– 200 lux cool white LED lighting. Experimental procedures approval from Duke Kunshan University Institutional Animal Care and Use Committee (project number: SKT-001) and were conducted in accordance with the Association for Assessment and Accreditation of Laboratory Animal Care (AAALAC) International standards (National Research Council, 2011).

### Operant chamber and light stimulus

Operant conditioning was conducted in a standard mouse operant chamber (16 cm × 14 cm × 13 cm; ENV-307A, Med Associates, Vermont) placed within a light-tight chamber (59.5 cm × 35.5 cm × 37.5 cm). The operant chamber had two short walls made of stainless steel and two long walls made of clear plastic (the front one served as the access door), with a clear plastic ceiling. Its floor consisted of a grid of 19 stainless-steel bars (0.3 cm diameter, spaced 0.5 cm apart), oriented parallel to the short walls; droppings fell through onto a tissue-lined tray beneath, which was cleaned between sessions. One short wall held a food receptacle (2.7 cm × 2.1 cm opening, 2 cm deep) positioned centrally and 0.2 cm above the grid floor; no food reward was used in this study. A stainless-steel response lever (1.6 cm × 0.8 cm paddle, 2.3 cm above the floor) was extended into the operant chamber at the start of the session and retracted at the end of the session. Reinforcement consisted of 4-s continuous illumination of a green LED light mounted 10.5 cm above the floor on the wall facing the receptacle. The emission spectrum of the green LED had a peak wavelength of 535 nm (measured using a spectrophotometer; XL-500 BLE, nanoLambda, Korea). Lever-press data were recorded and reinforcement schedules controlled by the Med-PC software (version V, Med Associates, Vermont).

### Intracranial virus injection

Mice were pre-assigned to two groups prior to operation: a control group (*n* = 6) receiving the control adeno-associated virus (AAV) vector carrying the enhanced green fluorescent protein (EGFP) alone (rAAV-hSyn-EGFP-WPRE-hGH-polyA, titer 5.17 × 10^12^ vg mL^−1^; catalogue number: PT-1990, Brain VTA, Wuhan, China); and an hM4Di group (*n* = 10) receiving the AAV vector carrying the inhibitory designer receptor hM4Di and the fluorescent tag [rAAV-hSyn-hM4D(Gi)-EGFP-WPRE-hGH-polyA, titer 5.07 × 10^12^ vg mL^−1^; catalogue number: PT-0153, Brain VTA, Wuhan, China]; the human synapsin (*hSyn*) promoter drives pan-neuronal viral expression at the injection site. The sample size of 6 mice in the control group is comparable to cohort sizes in previously published operant food-seeking studies (e.g., 4–8 mice per group in Shaw, Dawson, Reynolds, McCabe, & Leslie, 2004) and operant intracranial self-stimulation studies (e.g., 4–6 mice per group in Rossi, Sukharnikova, Hayrapetyan, Yang, & Yin, 2013). Pre-operation mean body weights were matched between control *versus* hM4Di groups: 28.917 ± 0.480 g *versus* 28.370 ± 0.332 g [*t*(14) = −0.966, *p* = 0.350].

During surgery, mice were anaesthetised with isoflurane, and the scalp was shaved, disinfected with iodine, and incised along the midline to expose the skull sutures. Bregma and lambda were identified and levelled (Z-axis difference <0.05 mm) by adjusting the incisor bar and ear bars, with bregma set as the zero coordinate. Burr holes were drilled bilaterally at anterior–posterior (AP) +0.50 mm and medial–lateral (ML) ±1.5 mm relative to bregma. The tip of the glass micropipette was lowered to dorsal–ventral (DV) −2.7 mm relative to dura, and AAV was infused via a microinjection pump at a rate of 0.05 µL min^−1^; a volume of 0.25 µL was delivered in each hemisphere. The micropipette was left *in situ* for 5 min after infusion before being slowly withdrawn from the injection site. The incision was rinsed with saline, sutured, and disinfected with iodine, after which mice received subcutaneous injection of meloxicam as analgesia. Mice recovered on a 37°C heating pad until fully ambulatory before being returned to their home cages. All mice fully recovered post-surgery and were included in subsequent behavioural training.

### Chemogenetic silencing during operant conditioning

Operant training began five weeks after surgery to ensure full recovery as well as sufficient AAV expression in the injection site. Thirty minutes prior to operant sessions, each mouse received intraperitoneal injection of 5 mg kg^−1^ clozapine N-oxide (CNO; catalogue number: 6329, Bio-Techne, Shanghai, China). This dose can activate hM4Di receptors and cause neuronal silencing *in vivo* in hM4Di-expressing mice for at least 3 hours (e.g., Milosavljevic, Cehajic-Kapetanovic, Procyk, & Lucas, 2016, see their Supplementary Figure *S*2C). The designer receptor agonist CNO was injected on days 1–3, day 5, and day 7 but was withheld on days 4 and 6 to assess non-specific effects of CNO in control mice (Traut et al., 2023) as well as the behavioural performance of hM4Di-expressing mice without the influence of CNO. Thirty minutes after CNO injection, the mouse was placed inside the operant chamber in darkness. At the start of the operant session, the response lever was inserted into the operant chamber; the left response lever was used as the operandum for half of the mice and the right response lever was used for the remaining animals. Lever-pressing responses were reinforced with 4-s green light delivered under fixed-ratio schedules: FR1 on days 1–2, FR3 on days 3–5, and FR5 on days 6–8. The operant chamber remained in darkness when no light reinforcer was being delivered. Each session was terminated when 30 light reinforcers had been delivered or when 45 min had elapsed since the start of the session, whichever occurred first. As light self-administration is more pronounced in the light phase than at night (Tam et al., 2026), mice received operant training between ZT01 and ZT04.

### Histology

Following completion of the behavioural experiment, mice were terminally anaesthetised with isoflurane and transcardially perfused with phosphate-buffered saline (PBS) followed by 4% paraformaldehyde (PFA). Brains were extracted, post-fixed in 4% PFA for 24 h, and cryoprotected in 30% sucrose for 24–48 h until fully submerged. Brains were extracted and immersed in 4% PFA for 24 h, then transferred to 30% sucrose solution for cryopreservation. After 2–3 days, brains were sectioned at 30 μm thickness with a cryostat. Brain sections were washed in PBS, stained with DAPI (1:3000) for 10 min, washed three times in PBS, and mounted on microscope slides. DAPI and GFP channels were imaged with a confocal microscope under a 10× objective lens.

### Statistical analyses

Means ± standard errors of the mean (SEM) were plotted in the figure, and statistical analyses were conducted in SPSS (version 31, IBM) and R (R Core Team, 2023); *α* = 0.05 (two-tailed) was adopted unless otherwise specified. First, to examine the relative contributions of Training Session, Lever Position (left *versus* right), Body Weight, Group, and Group × CNO interaction, linear mixed-effects models were conducted with these factors as fixed effects and individual mice as a random effect. The significance of each fixed effect was assessed using likelihood-ratio tests comparing the full model against a reduced model lacking the effect of interest (Bates, Mächler, Bolker, & Walker, 2015). A split-plot analysis of variance (ANOVA) was also conducted, with Training Session (days 1–8) as the within-subjects factor and Group (control *versus* hM4Di-expressing mice) as the between-subjects factor; Greenhouse–Geisser corrections were applied to the degrees of freedom (*df*) when the assumption of sphericity was violated; any *F* value with adjusted *df* was denoted as *FƐ*. Independent-samples *t* tests were used to compare lever-pressing responses between control and hM4Di groups during initial training (days 1–3) and to compare CNO effects between groups under FR3 and FR5 (days 4–8). One-sample *t* tests (two-tailed) were used to assess CNO effects against the value of zero.

## Results

In control and hM4Di mice, EGFP-positive somata are found in the dorsal striatum (caudate–putamen), confirming regional specificity of viral transfection to the dorsal but not ventral striatum (nucleus accumbens). Coronal brain sections showing EGFP immunofluorescence (*green*) in the caudate–putamen are displayed in **Fig. 1*a*,*b***. In addition, EGFP-positive projection fibres are found in the internal capsule, globus pallidus (GPe), and substantia nigra region (SNr), consistent with the known anatomical projections of dorsal striatal neurons in the mouse brain (see e.g., experiment `100142580` from Allen Mouse Brain Connectivity Atlas, Allen Institute for Brain Science, 2011; Oh et al., 2014); a sagittal brain section showing EGFP-positive projection fibres in the internal capsule, GPe, and SNr can be found in **Supplementary Fig. 1**. Injection of the designer receptor agonist CNO in control and hM4Di mice did not exert any noticeable non-specific effect on daily locomotor activity in home cages, which can be seen in the double-plotted actograms shown in **Fig. 1*c*,*d***.

**Fig. 1.**
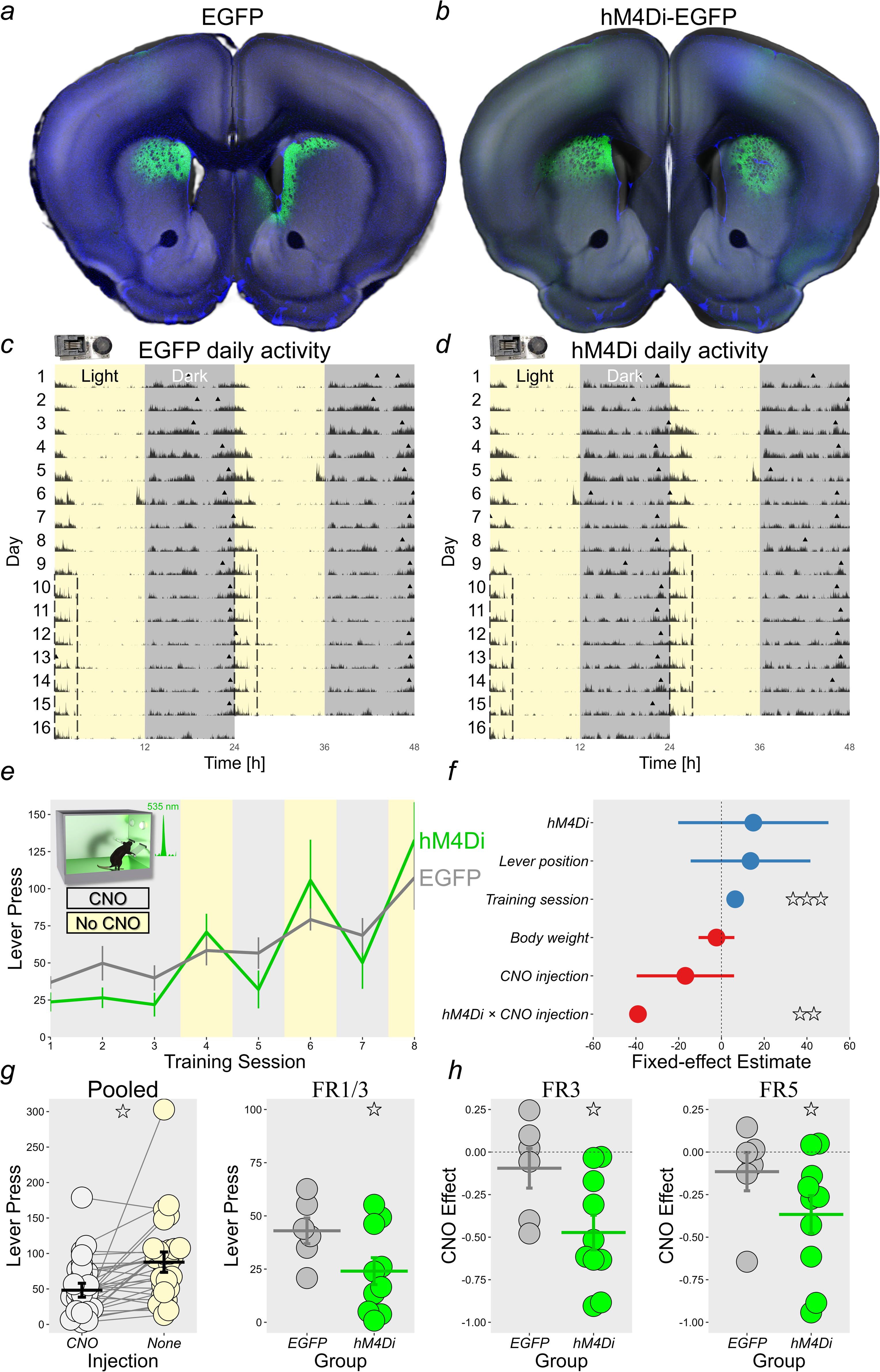
Effects of Dorsal Striatum Silencing on Light Self-administration in Mice. ***a*** and ***b*:** Coronal brain sections from control and hM4Di-expressing mice show the extent of adeno-associated virus (AAV) transfection in the dorsal striatum. The blue colour represents DAPI (nucleus) staining and the green colour represents enhanced green fluorescent protein (EGFP) expression. Brain sections were registered to cross sections in the Allen Mouse Brain 3D reference atlas (Wang et al., 2020) using DeepSlice (Carey et al., 2023). Cross-section images from the reference atlas in greyscale were overlaid on coronal brain sections using the QuickNII package (Neural Systems and Graphics Computing Laboratory, 2021). ***c*** and ***d*:** Double-plotted actograms show home cage locomotor activity monitored under passive-infrared (PIR) sensors (Brown et al., 2016). Each triangle indicates the PIR activity midpoint, which is defined as the time point at which activity in the preceding 8 h and activity in the subsequent 8 h are equated (Tam et al., 2021). Activity midpoints in both groups remained stable (i.e., not phase-shifted) before the experiment commenced and during the operant experiment (outlined by dashed rectangles). ***e*:** Lever presses in the operant experiment increased gradually in both groups (main effect of Session, *p* < 0.0001). ***f*:** Fixed-effect slope estimates with 95% confidence intervals from linear mixed-effects models confirmed the significance of Training Session (^⋆⋆⋆^*p* < 0.0001) and Group × CNO interaction (^⋆⋆^*p* = 0.00616). The forest plot was generated using the `sjPlot` package in R (Lüdecke et al., 2025). ***g*:** When pooled across groups, lever responding was generally higher on days when the chemogenetic actuator clozapine N-oxide (CNO) was withheld than when it was injected (^⋆^*p* = 0.0194, *left* panel); the effect of CNO remained as a non-significant trend (*p* = 0.0587) after the exclusion of the extreme data point with >300 lever presses from an hM4Di mouse tested under FR5 without CNO. On the first three sessions of operant training, hM4Di mice tended to respond less than control mice under the influence of CNO according to a one-tailed between-subjects *t* test (^⋆^*p* = 0.0322, *right* panel) with a large effect size (Cohen’s *d* = 1.037). ***h*:** Effects of CNO on FR3 and FR5 lever responding were quantified in terms of (*LNONE* − *LCNO*)/(*LNONE* + *LCNO*), where *LNONE* and *LCNO* represent lever presses on days without *versus* with CNO injection, respectively. CNO ratio scores under FR3 and FR5 were different from the value of zero in the hM4Di group [^⋆^*p*s = 0.00123 and 0.00907 (two-tailed), respectively], but this was not the case in the control group [*p*s = 0.453 and 0.352 (two-tailed), respectively]. After the exclusion of the hM4Di mouse with an extreme data point of >300 lever presses in panel ***g***, effects of dorsal striatal silencing by CNO in the hM4Di group remained significant under FR3 and FR5 [*p* = 0.00291 and 0.0172 (two-tailed), respectively].

Lever responding for light in the operant chambers increased gradually across the 8 days of training (**Fig. 1*e***). The linear mixed-effects model (**Fig. 1*f***) showed that there was an effect of Training Session [*χ*^2^(1) = 18.593, *p* < 0.0001], indicating acquisition of light self-administration under FR schedules as in our previous study (Tam et al., 2026). Crucially, there was a Group × CNO interaction [*χ*^2^(1) = 7.503, *p* = 0.00616], indicating that the designer receptor agonist CNO exerted a stronger detrimental effect on lever responding in hM4Di-expressing mice than in non-hM4Di control mice. The ANOVA confirmed the main effect of Training Session [Greenhouse–Geisser corrected *FƐ*(2,27) = 12.778, *p <* 0.0001, partial *η*^2^ = 0.516]. In the first three days of operant training, both control and hM4Di groups received systemic injection of 5 mg kg^−1^ CNO, a dose activating hM4Di receptors and causing neuronal silencing *in vivo* for at least 3 hours (Milosavljevic et al., 2016). Thirty minutes after CNO injection, hM4Di-expressing mice tended to show fewer lever-pressing responses than control mice, 24.036 ± 6.313 *versus* 43.002 ± 6.053 presses per 30 min, respectively, although this difference only reached significance according to a one-tailed between-subjects *t* test [*t*(14) = 2.007, *p* = 0.0322, Cohen’s *d* = 1.037]. This suggests that silencing of dorsal striatal neurons tended to attenuate lever responding during initial operant training sessions (**Fig. 1*g***).

To take into account potential non-specific effects of CNO injection (Traut et al., 2023), a ratio score was determined for each mouse: (*LNONE* − *LCNO*)/(*LNONE* + *LCNO*), where *LNONE* represents lever presses on days without CNO injection (days 4, 6, and 8) and *LCNO* represents lever presses under the influence of CNO (days 5 and 7); *ratio* ≈ 0 indicates negligible CNO effect and *ratio* < 0 indicates CNO-induced disruption of lever responding. In the control group, CNO had a negligible effect on lever presses [mean ± SEM under FR3: −0.095 ± 0.117; *t*(5) = −0.814, *p* = 0.453; mean ± SEM under FR5: −0.115 ± 0.112; *t*(5) = −1.026, *p* = 0.352; **Fig. 1*h***, *grey* points]. By contrast, lever responding in hM4Di mice was attenuated by CNO under FR3 and FR5 [mean ratio ± SEM under FR3: −0.473 ± 0.102; *t*(9) = − 4.634, *p* = 0.00123, Cohen’s *d* = −1.465; mean ratio ± SEM under FR5: −0.366 ± 0.111; *t*(5) = −3.311, *p* = 0.00907, Cohen’s *d* = −1.047; **Fig. 1*h***, *green* points]. The effect of Group on CNO ratio scores was significant under FR3 [*t*(14) = 2.359, *p* = 0.0334, Cohen’s *d* = 1.218] although not under FR5 [*t*(14) = 1.496, *p* = 0.157]. Taken together, chemogenetic silencing of dorsal striatal neurons transiently attenuated light self-administration but responding recovered on days when CNO was withheld, suggesting that behavioural performance rather than response acquisition was disrupted.

## Discussion

In the light self-administration paradigm (Tam et al., 2026), mice learn to lever-press to obtain a few seconds of light in the dark operant chamber. This acquired response is not driven by physiological needs such as hunger, as animals are not food or water deprived and no food reward is used during behavioural training. In addition, the response is not driven by novelty seeking, as the light reinforcer remains the same throughout training and thus reinforcer novelty should decrease rather than increase with continued training. This paradigm can be considered a simplified model relevant to digital technology-based disorders and behavioural addictions, offering an experimental approach to examine how behavioural persistence can arise due to light reinforcement contingency (Tam & Becker, 2026).

In the current study, chemogenetic silencing of dorsal striatal neurons in hM4Di-expressing mice attenuated light self-administration during initial days of training and under higher ratio schedules. However, response acquisition *per se* was not completely impaired, as lever responding recovered on days when the designer receptor agonist CNO was withheld. Crucially, CNO had a negligible effect on lever responding in the EGFP control group. This latter result suggests that the CNO effect in hM4Di-expressing mice was not due to potential reverse metabolism of CNO to clozapine, which is an atypical antipsychotic that crosses the blood–brain barrier more readily than CNO itself and binds to multiple endogenous (i.e., non-hM4Di) receptors (Gomez et al., 2017). Our results complement the established role of the dorsal striatum in food and drug self-administration in rodents (Everitt & Robbins, 2005, 2013; Yin & Knowlton, 2006) and extend its role to light self-administration. Combining the light self-administration paradigm with a loss-of-function manipulation (which is not feasible in human studies) allows us to reveal overlapping brain regions involved in light and drug self-administration in mice, bridging the gap between animal models and human neuroimaging studies reporting shared mechanisms underlying non-drug and drug habits (Klugah-Brown et al., 2021) and flanking a growing number of studies reporting associations between escalating progressive use of digital technology and increasing morphological and functional alterations in the dorsal striatum—in addition to the ventral striatum—e.g., in internet and online gaming disorders (Dong et al., 2021; Klugah-Brown et al., 2022; Yu et al., 2022).

While our findings demonstrate a causal role of the dorsal striatum in light self-administration in mice, converging evidence from human studies indicate broader psychological and behavioural effects of artificial light; for example, a recent review has summarised the effects of light on motivation and reward processing in humans (Mahoney & Schmidt, 2024, p. 163). In the context of behavioural addiction, although content-derived reinforcement is crucial in driving digital device use (File, 2026; Tunney, 2026), sensory reinforcement such as light emitted from LED screens also plays a role. Recent studies have reported that applying a greyscale filter on the smartphone—which removes colour while leaving all other functions intact on the device—can reduce screen time in young adults (Dekker & Baumgartner, 2023; Holte & Ferraro, 2020; Holte, Giesen, & Ferraro, 2023; Wickord & Quaiser-Pohl, 2023; Zimmermann & Sobolev, 2023). This emerging body of evidence indicates that coloured light is behaviourally rewarding, a notion that is consistent with the hypothesised hedonic property of light on other operandum-like devices, such as slot machines (Griffiths, 1993).

The hedonic property of light has been increasingly recognised in multiple psychiatric conditions (Huang, Tao, & Ren, 2024). For example, a recent meta-analysis of 11 randomised controlled trials (RCTs) with >800 patients has confirmed that bright light therapy (relative to dim red-light exposure) is effective in reducing depressive symptoms (Menegaz de Almeida et al., 2025). Light may have a similar therapeutic potential for behavioural addiction: a recent randomised controlled trial (Li et al., 2026) has reported that two weeks of bright light exposure (>5000 lux, 30 min per day) can reduce symptom severity and cue-induced craving in individuals with (Internet) gaming disorder relative to a dim-light condition (<200 lux). These improvements are accompanied by fMRI-measured functional connectivity changes in reward-related brain regions (Li et al., 2026). Thus, light is a “double-edged sword” as it can reshape brain reward circuits, potentially alleviating addictive symptoms (Li et al., 2026) and improving mood in certain clinical populations (Menegaz de Almeida et al., 2025), but on the other hand paired with reinforcement-based algorithms and personalised content may shape maladaptive digital habits in vulnerable individuals (Dekker & Baumgartner, 2023; Holte & Ferraro, 2020; Holte et al., 2023; Wickord & Quaiser-Pohl, 2023; Zimmermann & Sobolev, 2023). The mechanisms of these different effects of light and its contribution to addictive behaviour remain to be fully understood (Huang et al., 2024). Together with our previous empirical findings (Tam et al., 2026), the initial rewarding experience of response-contingent light delivered under traditional reinforcement schedules can give rise to behavioural persistence that is partly driven by the dorsal striatum, demonstrating overlapping neural mechanisms between non-drug and drug habits in laboratory rodents and potentially between certain behavioural addictions such as digital technology-based disorders and drug addiction in humans.

Regarding the limitations of the current study, it remains to be determined what specific processes are affected by dorsal striatal silencing. Under higher ratio schedules (FR3 and FR5), attenuated light self-administration in CNO-treated hM4Di-expressing mice could be due to reduced sensitivity to the hedonic value of light or reduced habitual responding, or a combination thereof. Future work can dissociate these possibilities by using the breaking point measure under a progressive-ratio schedule to assess the motivational significance of light (Richardson & Roberts, 1996) and adopting reinforcer-devaluation or contingency-degradation procedures to assess habitual control (Dickinson & Balleine, 1994). In addition, using multiple groups of striatal hM4Di-expressing mice receiving the designer receptor agonist in different phases (e.g., during the acquisition or extinction phase) would allow the time course of acquisition and extinction of light self-administration to be examined. Another limitation of the current study is the use of the pan-neuronal promoter human synapsin (*hSyn*) to drive expression of designer receptor hM4Di in the dorsal striatum, which precludes understanding the functional roles of different striatal cell types. Future investigations using cell-type-specific promoters can dissociate the contributions of direct D1-receptor-expressing and indirect D2-receptor-expressing striatal pathways (e.g., Kravitz, Tye, & Kreitzer, 2012) as well as different types of striatal interneurons (Yin, 2024), thereby providing a more complete understanding of the mechanisms underlying behavioural persistence towards light reinforcers. Finally, it should be noted that while light possesses some rewarding properties in both nocturnal mice and diurnal humans, species differences in light sensitivity and behavioural responses to light (Foster, Hughes, & Peirson, 2020) must be carefully considered to fully evaluate the potential translational relevance of the mouse light self-administration paradigm (File, 2026; Tunney, 2026).

---

**Supplementary Fig. 1. EGFP-positive Projection Fibre Distribution**

***a*:** In mice receiving AAV injection into the dorsal striatum (CP), EGFP+ projection fibres were found in the internal capsule (int), globus pallidus (GPe), and midbrain substantia nigra region (SNr); the blue colour represents DAPI (nucleus) staining and the green colour represents EGFP immunofluorescence on the sagittal brain section. ***b*:** Our result in panel ***a*** is consistent with the known anatomical projections of dorsal striatal neurons in the mouse brain, which can be seen from the results of the AAV injection experiment `100142580` in the Allen Mouse Brain Connectivity Atlas (Allen Institute for Brain Science, 2011; Oh et al., 2014). Supplementary Fig. 1 can be accessed via the following link: https://doi.org/10.6084/m9.figshare.33130181

